# Activation of the NAD⁺–Sirtuin Axis Protects Against Chronic Doxorubicin-Induced Subclinical Renal Tubular Injury Through Restoration of Mitochondrial Homeostasis and Suppression of Inflammation

**DOI:** 10.64898/2026.07.29.741470

**Authors:** Keiji Saito, Ryusuke Hosoda, Ryo Numazawa, Tomoki Hori, Iyori Nojima, Yukika Saga, Yuki Tatekoshi, Tatsuya Sato, Koki Abe, Atsushi Kuno

## Abstract

**Background and purpose:** Anthracyclines, such as doxorubicin (DOX), are associated with late-onset kidney dysfunction; however, the mechanisms underlying chronic tubular injury remain poorly understood. We investigated whether chronic low-dose DOX exposure induces persistent mitochondrial dysfunction in renal tubules and evaluated the therapeutic potential of activating the NAD⁺–Sirtuin axis.

**Experimental Approach:** C57BL/6 mice were repeatedly administered low-dose DOX with or without resveratrol (RSV) or nicotinamide mononucleotide (NMN), a sirtuin activator. Renal injury was assessed using neutrophil gelatinase-associated lipocalin (NGAL) staining. Integrated proteomic and RNA sequencing analyses were performed to identify molecular alterations. Mitochondrial morphology and function were evaluated using structured illumination microscopy (SIM) of Masson’s trichrome-stained paraffin sections and *ex vivo* Seahorse analysis of freshly isolated renal tubules.

**Key Results:** Chronic DOX administration induced tubular injury, despite preserving serum creatinine levels. Multi-omics analyses consistently demonstrated the suppression of mitochondrial pathways, including oxidative phosphorylation, fatty acid oxidation, and mitochondrial gene expression. SIM revealed mitochondrial fragmentation in tubular epithelial cells, whereas the Seahorse assay showed impaired mitochondrial respiratory capacity in isolated renal tubules. DOX also increased tubular acetylated superoxide dismutase 2 (SOD2) levels and activated inflammatory pathways. Importantly, both RSV and NMN attenuated tubular injury, restored mitochondrial metabolic pathways, reduced SOD2 acetylation, improved mitochondrial morphology, and suppressed inflammatory responses.

**Conclusions and Implications:** Chronic low-dose DOX exposure induces subclinical renal tubular injury characterized by mitochondrial dysfunction and inflammation. The pharmacological activation of sirtuins confers reno-protective effects by preserving mitochondrial homeostasis. These findings identify mitochondrial dysfunction as a central therapeutic target in DOX-induced nephrotoxicity and support sirtuin modulation as a potential strategy for preventing chemotherapy-related chronic kidney injury.

**Bullet point summary:** *What is already known:*

- Doxorubicin causes cardiotoxicity through mitochondrial dysfunction and oxidative stress.
- Doxorubicin-induced tubular injury and the associated late-onset kidney dysfunction are clinically proven.

*What does this study add:*

- Chronic low-dose doxorubicin induces tubular mitochondrial dysfunction, as identified by integrated multi-omics analyses.
- Resveratrol and NMN preserve mitochondrial integrity and suppress inflammatory responses in renal tubules.

*Clinical significance:*

- Mitochondrial dysfunction may represent an early therapeutic target in doxorubicin-associated nephrotoxicity.
- Activation of the NAD⁺–Sirtuin axis could prevent chronic kidney injury in cancer survivors.

## Introduction

Advances in cancer therapy have markedly improved long-term survival, shifting clinical priorities toward the management of treatment-related complications (Kooijmans et al. 2025). Among these, late-onset chronic kidney disease (CKD) has emerged as a major determinant of morbidity and quality of life in cancer survivors. Recent clinical evidence suggests that anthracyclines, such as doxorubicin (DOX), in addition to their well-recognized cardiotoxicity, may also contribute to renal tubular injury and progressive chronic renal dysfunction (Dieffenbach et al. 2021; Bárdi et al. 2007). Thus, elucidating the mechanisms underlying DOX-induced tubular injury and developing targeted therapeutic strategies to prevent and treat this condition are urgently warranted. However, most experimental studies have relied on high-dose acute injury models that do not recapitulate chronic renal impairment observed after repeated low-dose chemotherapy in cancer survivors and patients receiving long-term chemotherapy (Lee and Harris 2011; Zheng et al. 2006, 2005). Consequently, the mechanisms by which chronic DOX exposure drives CKD progression remain poorly understood, thereby limiting the development of targeted preventive strategies.

DOX-induced cardiomyopathy is a well-recognized clinical complication and is mechanistically linked to DNA damage, mitochondrial dysfunction, cell death, and progressive inflammation and fibrosis (Wu et al. 2024; Li et al. 2019; Kuno et al. 2023; Wallace, Sardão, and Oliveira 2020). DOX accumulates within the mitochondria in cardiomyocytes, causing mitochondrial DNA depletion, impaired transcription, reduced mitochondrial mass, and the suppression of fatty acid oxidation, ultimately leading to sustained energy failure. Similar to cardiomyocytes, renal tubular epithelial cells are highly enriched in mitochondria due to the substantial energy demands of active transport and metabolic homeostasis (Doke and Susztak 2022; Brown et al. 2017). These observations suggest that mitochondrial dysfunction may represent a central pathogenic mechanism in chronic DOX-induced renal injury.

Sirtuins are NAD⁺-dependent deacetylases that regulate mitochondrial homeostasis and cellular metabolism. Among the sirtuin family, Sirtuin-1 (SIRT1) and Sirtuin-3 (SIRT3) have been most consistently reported to exert protective effects against kidney injury. Studies using knockout mouse models have demonstrated that deficiency in either SIRT1 or SIRT3 exacerbates renal ischemia-reperfusion injury (Cheng et al. 2022; Wakino, Hasegawa, and Itoh 2015; Fan et al. 2013). Mechanistically, SIRT1 promotes mitochondrial biogenesis, whereas SIRT3 primarily maintains mitochondrial homeostasis and antioxidant defense, including activation of superoxide dismutase 2 (SOD2). Recent reports have shown that the pharmacological activation of SIRT1 by resveratrol (RSV) attenuates DOX-induced cardiomyopathy, supporting the protective role of sirtuins against DOX-induced mitochondrial injury (Danz et al. 2009). Nicotinamide mononucleotide (NMN) has also been reported to activate SIRT1 signaling by restoring intracellular NAD⁺ levels, thereby improving mitochondrial function and protecting against tissue injury (Yamamoto et al. 2014). However, it remains unclear whether RSV or NMN attenuates DOX-induced tubular injury.

In this study, we established a clinically relevant chronic low-dose DOX mouse model to investigate the mechanisms underlying anthracycline-induced tubular injury. By integrating proteomic and transcriptomic analyses with complementary structural and functional assessments of mitochondria, we identified renal tubular mitochondrial dysfunction as the central pathogenic mechanism of chronic DOX nephrotoxicity. Furthermore, we demonstrated structural and functional mitochondrial impairments using structured illumination microscopy of paraffin-embedded kidney sections and Seahorse metabolic flux analysis of freshly isolated renal tubules. Finally, we showed that pharmacological activation of the NAD⁺–Sirtuin axis by RSV or NMN ameliorates mitochondrial dysfunction and protects against chronic DOX-induced tubular injury.

## Methods

### 2.1. Animals

All animal experiments were performed as described previously (Kuno et al. 2023). All procedures were performed according to the Animal Guidelines of Sapporo Medical University and were approved by the Animal Use Committee (#25-018). This study was performed in accordance with the ARRIVE guidelines and conformed to the Guide for the Care and Use of Laboratory Animals published by the US National Institutes of Health (NIH Publication, 8th Edition, 2011).

To establish a DOX-induced nephrotoxicity model, male C57BL/6N mice were randomly divided into four groups: Vehicle, DOX, RSV + DOX, and NMN + DOX. As reported previously (Kuno et al. 2023), mice in the Vehicle and DOX groups received vehicle (phosphate buffer saline [PBS]) and DOX (LC Laboratories, Woburn, MA, USA; 5 mg/kg in PBS), respectively, via intraperitoneal (i.p.) injection once a week for four weeks starting at 13 weeks of age. Mice in the RSV+DOX group were administered RSV (R0071, Tokyo Chemical Industry, Tokyo, Japan; 0.4 g/kg food) ad libitum from 12 weeks of age and received DOX as in the DOX group. Mice in the NMN+DOX group were administered NMN (Oriental Yeast Co., Ltd., Tokyo, Japan; 0.5 g/kg i.p.) 30 min before and 2 days after each DOX injection, and received DOX as in the DOX group. The RSV dose was determined based on a previous report (Kuno et al. 2023). One week after the final DOX injection, the mice were euthanized under deep anesthesia with 5% isoflurane, followed by cervical dislocation. Serum and kidney tissues were collected for subsequent analyses (Fig. 1A).

**Figure 1.**
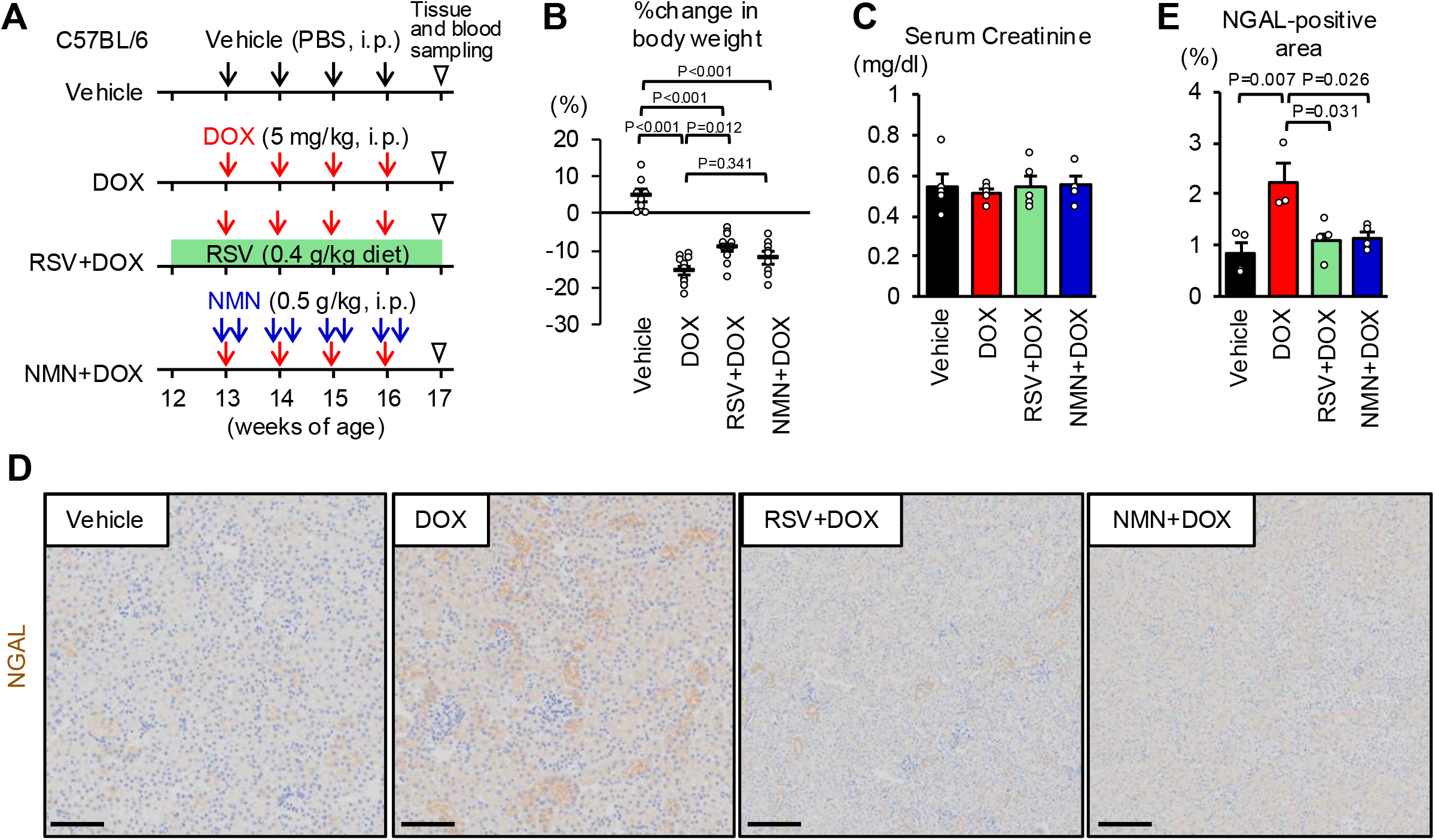
Chronic doxorubicin (DOX) administration induces subclinical tubular injury, which is attenuated by resveratrol (RSV) and nicotinamide mononucleotide (NMN). (A) Experimental protocol. Mice received intraperitoneal injections of vehicle (phosphate buffer saline [PBS]) or doxorubicin (DOX, 5 mg/kg) once weekly for four weeks, starting at 13 weeks of age. Mice in the RSV+DOX group were fed an RSV-containing diet (0.4 g/kg diet) beginning 1 week before the first DOX injection until sacrifice. Mice in the NMN+DOX group received NMN (0.5 g/kg) intraperitoneally, 30 min before and 2 days after each DOX injection. Kidney tissues and blood samples were collected one week after the final DOX injection. (B) Changes in body weight after doxorubicin treatment. N=8–10 per group. (C) Serum creatinine levels one week after the final DOX injection. N=4–5 per group. (D) Representative images of neutrophil gelatinase-associated lipocalin (NGAL) immunostaining. Scale bar: 100 µm. (E) Quantification of NGAL-positive areas. N=3–4 per group. Data are presented as mean ± SE. Statistical analysis was performed using a one-way analysis of variance (ANOVA) followed by Tukey’s multiple comparison test.

### 2.2. Proteomic analysis

For each sample, 10 μg of protein lysate was reduced with 10 mM dithiothreitol at 56 ℃ for 30 min, then alkylated with 27.5 mM iodoacetamide in the dark at room temperature for 30 min. Protein purification and digestion were performed using single-pot solid-phase-enhanced sample preparation (SP3) with minor modifications (Hughes et al. 2019). Tryptic digestion was performed using 500 ng Trypsin/Lys-C Mix (Promega, Madison, WI, USA) overnight at 37 ℃. The digested peptides were desalted using a GL-Tip SDB (GL Sciences, Tokyo, Japan) according to the manufacturer’s instructions and dried in a centrifugal evaporator. The dried peptides were reconstituted in 2% acetonitrile (ACN) containing 0.1% formic acid (FA).

The reconstituted peptides were loaded into a 75 μm × 12 cm capillary column packed with 3 μm particles (Nikkyo Technos, Tokyo, Japan) and separated using an EASY-nLC 1000 system (Thermo Fisher Scientific) at a flow rate of 300 nL/min. Mobile phase A comprised 0.1% FA in water, and mobile phase B comprised 0.1% FA in ACN. Peptides were separated using a 105 min gradient: 5–35% B over 90 min, 35–95% B over 5 min, and a final hold at 95% B for 10 min. Peptides eluted from the column were analyzed on a Q Exactive Plus Orbitrap mass spectrometer (Thermo Fisher Scientific) in the data-independent acquisition (DIA) mode. Full MS spectra were acquired over the m/z 500–860 range at a resolution of 35,000, with an automatic gain control (AGC) target of 3 × 10^6^ and a maximum injection time of 110 ms. Sixty consecutive isolation windows of 6 m/z were used to fragment all precursor ions from m/z 500 to 860 per DIA cycle. The MS/MS spectra were acquired at a resolution of 17,500 with an AGC target of 1 × 106 and a maximum injection time of 50 ms.

Raw DIA data were processed using DIA-NN (version 2.3.0) against an in silico spectral library generated from the UniProt reference proteome of Mus musculus (Proteome ID: UP000000589; 17,218 reviewed proteins). The spectral library was predicted from the FASTA sequences within DIA-NN using the following parameters: trypsin/P as the protease with a maximum of one missed cleavage, protein N-terminal methionine excision enabled, carbamidomethylation of cysteine as a fixed modification, no variable modifications, peptide length of 7–45 amino acids, precursor charge of 2–4, precursor m/z from 500 to 860, and fragment ion m/z from 200 to 1800. For the DIA-NN search, the following parameters were used: mass accuracy at MS1 and MS2 was set to 10Oppm, the scan window was set to 6, match-between-runs and protein inference were enabled, scoring was set to generic, machine learning was set to NNs (cross-validated), quantification strategy was set to QuantUMS (precision), and cross-run normalization was set to RT-dependent. The “--peak-translation” command was applied in the DIA-NN command line. Protein identification was filtered at a 1% false discovery rate at both the precursor and protein levels. Protein quantification was performed using the intensity matrix generated by DIA-NN.

Datasets with an overall missing-value rate greater than 30% were excluded from the analysis. In addition, only the proteins detected in at least 50% of the samples within each group were retained. Missing values were imputed separately within each group: K-nearest neighbor (KNN) imputation was applied when the within-group missing value fraction was ≤20%, whereas quantile regression imputation of left-censored data (QRILC) was applied when the within-group missing value fraction was >20%.

Differential protein expression analysis was performed using the limma package. For each comparison, linear models were fitted using groupwise matrices, followed by empirical Bayes moderation.

Gene set enrichment analysis (GSEA) was performed using the ClusterProfiler and ReactomePA packages. Proteins were ranked according to moderated t-statistics from the differential analysis. Enrichment analyses were performed for Gene Ontology biological processes, molecular functions, and cellular components, as well as Reactome pathways.

### 2.3. Gene expression analysis using RNA-seq

Total RNA was isolated from the kidneys as described previously (Kimura et al. 2019). The mRNA samples were processed and sequenced by Rhelixa Co., Ltd., Tokyo, Japan. Datasets with an overall missing-value rate greater than 30% were excluded from the analysis. In addition, only the proteins detected in at least 50% of the samples within each group were retained. Missing values were imputed separately within each group: KNN imputation was applied when the within-group missing value fraction was ≤20%, whereas QRILC was applied when the within-group missing fraction was >20%.

Differential protein expression analysis was performed using the limma package. For each comparison, linear models were fitted using groupwise matrices, followed by empirical Bayes moderation.

GSEA was performed using ClusterProfiler and ReactomePA. Proteins were ranked according to moderated t-statistics from the differential analysis. Enrichment analyses were performed for Gene Ontology biological processes, molecular functions, cellular components, and Reactome pathways.

### 2.4. Immunohistochemistry

Immunohistochemistry was performed as described previously (Kimura et al. 2019). After fixation with 10% formalin, paraffin-embedded kidney samples were cut into 3 μm sections. Kidney sections were stained with primary antibodies against neutrophil gelatinase-associated lipocalin (NGAL; ab216462, 1:100; Abcam, Cambridge, UK) and acetyl (K68)-SOD2 (AcSOD2; ab137037, 1:250) at the Biomedical Research Center Division of Morphological Research, Sapporo Medical University. NGAL-positive areas were identified in five randomly selected fields from three to four kidneys per group. The glomerular area was excluded from the NGAL analysis. The Ac-SOD2-positive signal intensity was quantified, and the mean intensity of each field was measured in four randomly selected fields per mouse, from seven to eight mice per group.

### 2.5. **Analysis of mitochondrial morphology by** structured illumination microscopy (**SIM) of Masson-Trichrome staining**

The mitochondrial morphology was evaluated using previously published methods (Matsumoto et al. 2021). Formalin-fixed, paraffin-embedded kidney tissues were sectioned and stained with Masson’s trichrome. Mitochondrial morphology in renal tubular cells was further assessed using SIM under standardized imaging conditions. Kidneys from four mice per group were examined, and three to four representative cortical fields were acquired from each kidney under identical imaging conditions. For the morphometric analysis, four tubular cells per group were randomly selected and subjected to mitochondrial segmentation analysis. The major and minor axes of each mitochondrion were measured, and the mitochondrial aspect ratio was calculated as the ratio of major axis length to minor axis length. Approximately 2,000–3,000 mitochondria pooled from all the cells per group were included in the analysis.

### 2.6. Measurement of mitochondrial respiration of freshly isolated tubular cells

Proximal tubular segments were isolated from the mouse renal cortex and subjected to bioenergetic analysis using the Seahorse XF platform (Ogawa et al. 2023; Ichimura et al. 2008; Hammoud et al. 2025). Vehicle- or DOX mice (8 mg/kg i.p. once a week for four weeks) were euthanized, and the kidneys were rapidly excised. Renal cortices were dissected, finely minced, and digested in HBSS containing collagenase type IV (590-02793, MP Biomedicals, Irvine, CA, USA; 1 mg/mL) at 37 ℃ for 30 minutes with gentle agitation. The reaction was stopped by adding ice-cold PBS, and the suspension was mechanically dispersed. The glomeruli and the remaining tissue clumps were separated by gravity sedimentation for 1 min. The supernatant containing the renal tubules was passed through a 100-µm cell strainer, and the filtrate was centrifuged at 300 × g for 5 min. The resulting pellet was collected, washed twice with PBS, and resuspended in Seahorse XF DMEM (pH 7.4; Agilent Technologies, #103575-100), supplemented with 5.5 mM glucose, 2.0 mM glutamine, and 1.0 mM sodium pyruvate. An aliquot of the tubule suspension was lysed using CelLytic MT Cell Lysis Reagent (Sigma-Aldrich, #C3228), and the protein concentration was determined using a BCA assay (TaKaRa Bio, #T9300A). Approximately 15 μg of protein-equivalent tubular suspension was seeded into each well of a Seahorse XFe96 Cell Culture Microplate (Agilent Technologies, #103794-100) pre-coated with Cell-Tak (Corning, #354240), based on preliminary optimization of the assay conditions. The plates were then equilibrated in a COO-free incubator at 37 ℃ for 1 h, and the final volume in each well was adjusted to 180 μL with assay medium immediately before measurement. The oxygen consumption rate (OCR) was measured using a Seahorse XFe96 Bioanalyzer (Agilent Technologies) according to the manufacturer’s instructions. OCR was assessed at baseline and after sequential injections of oligomycin (final concentration, 1.0 μM), carbonyl cyanide p-trifluoromethoxyphenylhydrazone (FCCP; final concentration, 5.0 μM), and a mixture of rotenone and antimycin A (final concentration, 1.0 μM each). Each measurement cycle comprised 3 min of mixing followed by 3 min of measurement. OCR values were normalized to protein content to correct for variability in the amount of seeded tubular material among the wells.

### 2.7. Measurements of serum creatinine

Serum creatinine levels were measured using a QuantiChrom Creatinine Assay Kit (DICT-500, BioAssay Systems, CA, USA).

### 2.8. Statistical analysis

SigmaPlot software (version 14.0) and GraphPad Prism software were used for statistical analyses. Data are shown as mean ± SE. The unpaired 2-tailed Student’s t-test was used to compare two groups. Statistical significance between more than three groups was evaluated using parametric and nonparametric one-way analyses of variance. Proteomic and RNA-seq analyses were conducted using R version 4.3.0 (R Foundation for Statistical Computing, Vienna, Austria). Tukey’s test was used for multiple comparisons. Statistical significance was set at *P* < 0.05.

## Results

### 3.1. Both NMN and RSV Independently Attenuate Tubular Injury Induced by Chronic Low-Dose DOX

To investigate the mechanisms underlying chronic DOX-induced renal injury, we established a chronic low-dose DOX administration model (Fig. 1A). C57BL/6N mice received DOX (5 mg/kg) once weekly for four consecutive weeks. To evaluate therapeutic potential, additional groups were treated with the SIRT1 activator RSV or the NAD⁺ precursor NMN in combination with DOX. The kidneys were harvested five weeks after the initial injection. Chronic DOX administration was well tolerated, with no observed acute mortality. Although DOX-treated mice exhibited alterations in body weight (Fig. 1B), serum creatinine levels remained unchanged compared with those in the Vehicle group (Fig. 1C), indicating preserved global renal function.

Because drug-induced nephrotoxicity may initially manifest as tubular injury prior to detectable changes in serum creatinine, we assessed tubular damage using NGAL immunostaining. DOX treatment significantly increased the NGAL-positive tubular area, demonstrating tubular injury despite preserved renal function (Fig. 1D, E). Importantly, RSV significantly attenuated DOX-induced body-weight loss, whereas NMN showed a similar but non-significant trend. In contrast, both RSV and NMN significantly reduced the NGAL-positive area without affecting the serum creatinine levels (Fig. 1B–E).

Together, these findings establish a clinically relevant model of chronic low-dose DOX-induced tubular injury and demonstrate that pharmacological activation of the sirtuin pathway mitigates DOX-associated tubular damage.

### 3.2. Both RSV and NMN Suppress DOX-Induced Acetylation of SOD2 in Renal Tubules

As RSV and NMN are known to promote deacetylation by activating SIRT1 and SIRT3, we examined mitochondrial protein acetylation by assessing acetylated SOD2 levels in renal tubules. Immunostaining revealed that chronic DOX administration significantly increased SOD2 acetylation in the tubular epithelial cells (Fig. 2A, B). Importantly, the co-administration of RSV or NMN markedly attenuated DOX-induced SOD2 acetylation in the renal tubules (Fig. 2A, B). These findings indicate that RSV and NMN activate the deacetylation pathways in tubular cells under chronic DOX exposure.

**Figure 2.**
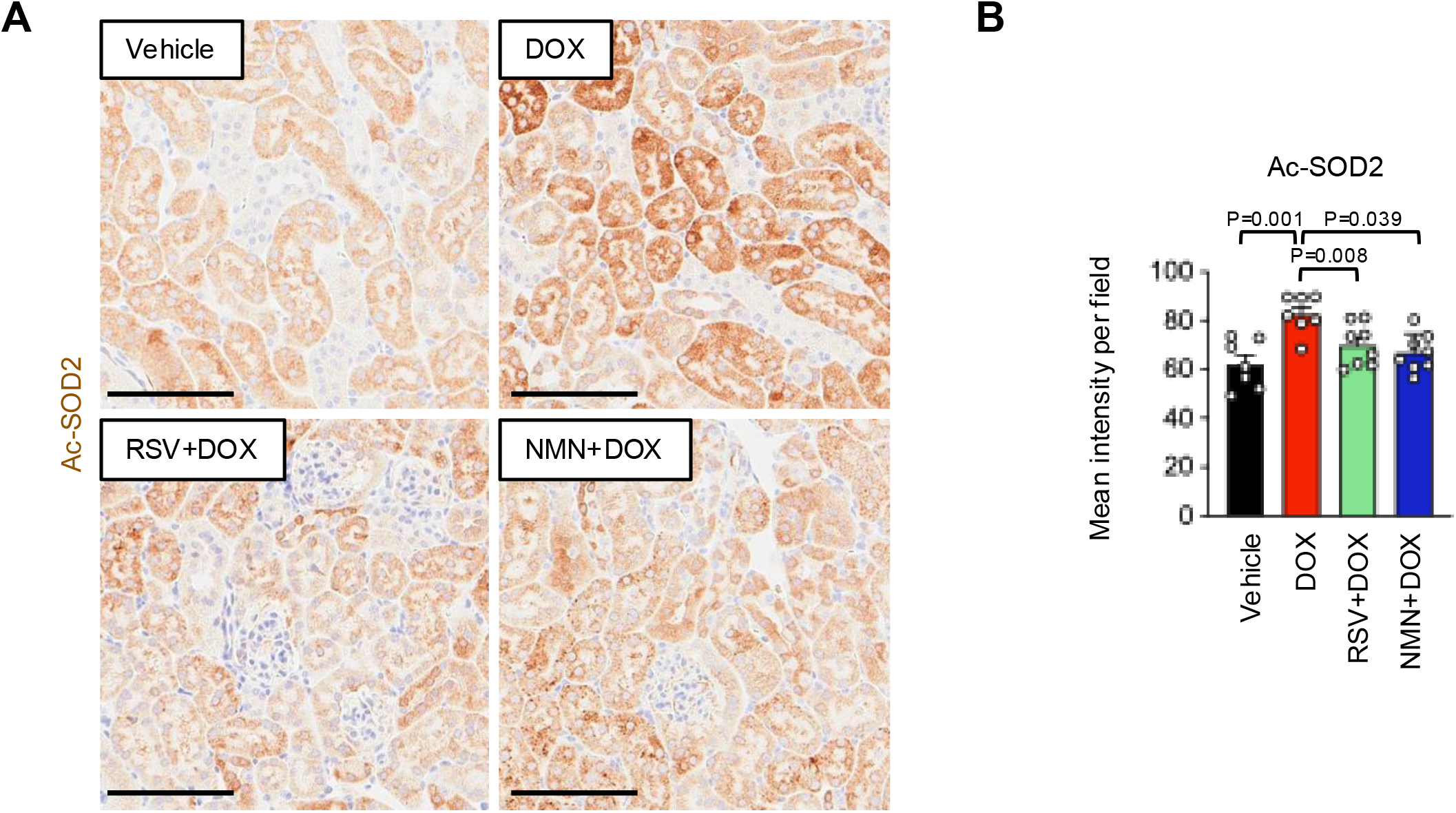
RSV and NMN suppress DOX-induced acetylation of mitochondrial protein. (A) Representative images of acetylated superoxide dismutase 2 (Ac-SOD2) immunostaining. Scale bar 100 µm. (B) Quantification of Ac-SOD2 mean intensity. N=7–8 per group. Statistical analysis was performed using a one-way ANOVA followed by Tukey’s multiple comparison test.

### 3.3. Multi-Omics Analyses Reveal Mitochondrial Dysfunction Induced by Chronic DOX

To elucidate the molecular mechanisms underlying DOX-induced tubular injury and the protective effects of RSV and NMN, we performed integrated proteomic and RNA sequencing analyses using renal tissues from the four groups: Vehicle, DOX, RSV+DOX, and NMN+DOX (Fig. 3A).

**Figure 3.**
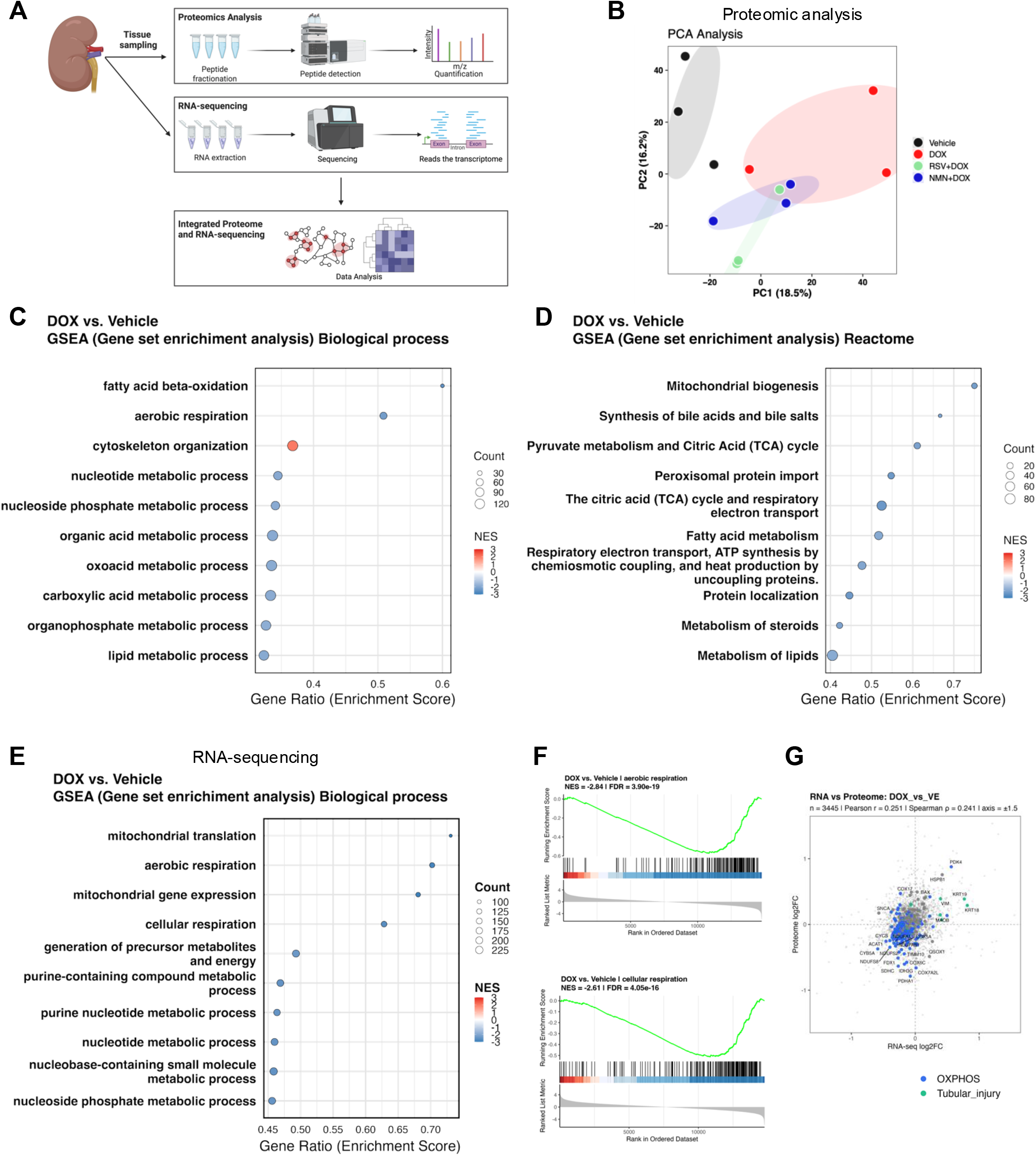
Integrated transcriptomic and proteomic analyses identify mitochondrial dysfunction following chronic DOX treatment. (A) Overview of the experimental workflow. Kidney tissues underwent proteomic analysis and RNA sequencing, followed by integrated multi-omics data analysis. (B) Principal component analysis of proteomic profiles. (C) Gene set enrichment analysis (GSEA) of proteomic data comparing DOX and Vehicle groups, showing enriched Gene Ontology biological process terms. (D) Reactome pathway enrichment analysis of proteomic data comparing DOX and Vehicle groups. (E) GSEA of RNA-seq data comparing DOX and Vehicle, showing enriched Gene Ontology biological process terms. (F) Representative GSEA enrichment plots for aerobic and cellular respiration. (G) Integrated analysis of transcriptomic and proteomic alterations induced by DOX. Genes and proteins associated with oxidative phosphorylation and tubular injury are highlighted.

In the proteomic analysis, after standard preprocessing and quality control, a total of 3,664 proteins were identified and quantified. Principal component analysis (PCA) of protein expression profiles distinguished the DOX and Vehicle groups. Notably, the RSV+DOX and NMN+DOX groups clustered away from the DOX group and were closer to the controls, which was consistent with their phenotypic protection against tubular injury (Fig. 3B). To characterize the pathways altered by DOX exposure, we performed GSEA to compare the DOX and Vehicle groups. Proteomic data revealed significant downregulation of pathways related to fatty acid β-oxidation, mitochondrial biogenesis, and the tricarboxylic acid (TCA) cycle in DOX-treated kidneys (Fig. 3C, D).

We performed RNA sequencing to validate these findings at the transcriptomic level. Similarly, GSEA demonstrated significant suppression of pathways associated with mitochondrial translation, aerobic respiration, and mitochondrial gene expression in the DOX group compared with the Vehicle group (Fig. 3E, F). The enrichment scores were significantly altered in a direction consistent with mitochondrial dysfunction. Together, these proteomic and transcriptomic data consistently indicate that chronic DOX administration induces mitochondrial impairment in renal tissues. To assess the concordance between the transcriptomic and proteomic alterations induced by DOX, fold-changes relative to the Vehicle group were plotted for each gene (Fig. 3G). Notably, tubular injury-associated genes were clustered in the upper-right quadrant, indicating coordinated upregulation at both mRNA and protein levels. In contrast, genes involved in oxidative phosphorylation were predominantly localized in the lower-left quadrant, reflecting a consistent downregulation across both datasets. These findings demonstrate a strong agreement between the transcriptomic and proteomic analyses and support the robustness of the multi-omics approach in capturing biologically relevant changes induced by DOX.

### 3.5. RSV and NMN Restore Mitochondrial Pathways in Proteomic Analysis

To further delineate the mechanisms underlying tubular protection by RSV and NMN, we performed comparative GSEA between the DOX and RSV+DOX or NMN+DOX groups in the proteomic dataset. Both RSV and NMN treatments significantly reversed the suppression of mitochondria-related pathways observed in DOX-treated kidneys, including aerobic respiration, the TCA cycle, and mitochondrial biogenesis, indicating partial restoration of mitochondrial metabolic programs (Fig. 4A, B, C, D).

**Figure 4.**
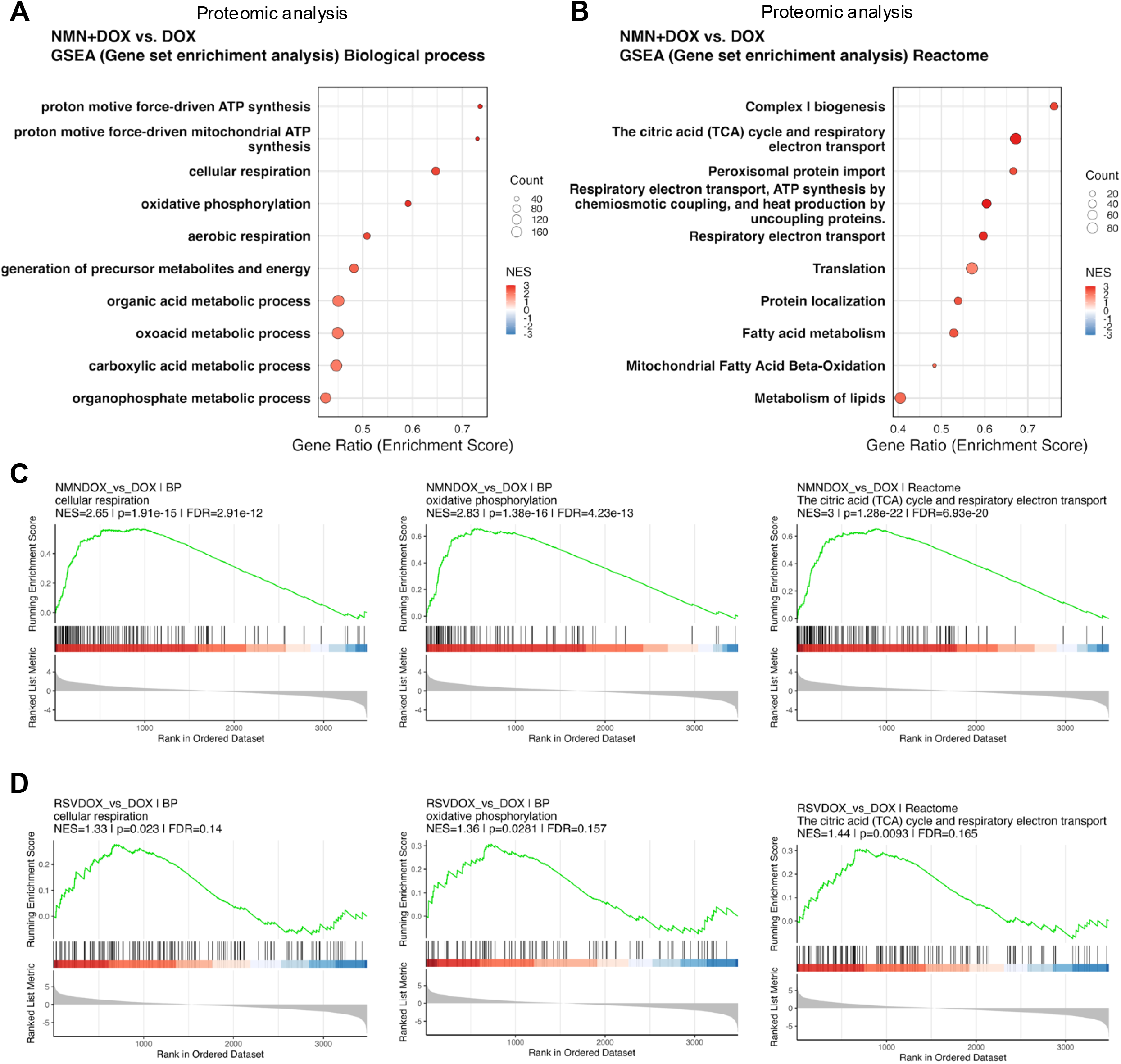
NMN and RSV restore mitochondrial pathways suppressed by chronic DOX treatment. (A) GSEA of proteomic data comparing NMN+DOX and DOX groups, showing enriched Gene Ontology biological process terms. (B) Reactome pathway enrichment analysis comparing the NMN+DOX and DOX groups. (C) Representative GSEA enrichment plots comparing the NMN+DOX and DOX groups. (D) Representative GSEA enrichment plots comparing the RSV+DOX and DOX groups.

### 3.6. Both RSV and NMN Attenuate DOX-Induced Mitochondrial Fragmentation in Renal Tubules

Heatmap analysis demonstrated that DOX treatment downregulated a broad range of genes encoding components of mitochondrial respiratory chain complexes, whereas additional NMN administration partially restored their expression. These findings suggest that DOX impairs mitochondrial integrity and function, and that NMN mitigates this mitochondrial dysfunction (Fig. 5A). To validate the mitochondrial alterations suggested by our multi-omics analyses, we directly assessed the mitochondrial morphology in renal tissue. Recent studies have demonstrated that SIM applied to paraffin-embedded Masson’s trichrome-stained kidney sections enables high-resolution visualization of mitochondrial structures without the need for electron microscopy (Matsumoto et al. 2021). Using this approach, the mitochondrial morphology was clearly visualized in the renal tubules of a chronic DOX nephropathy model (Fig. 5B).

**Figure 5.**
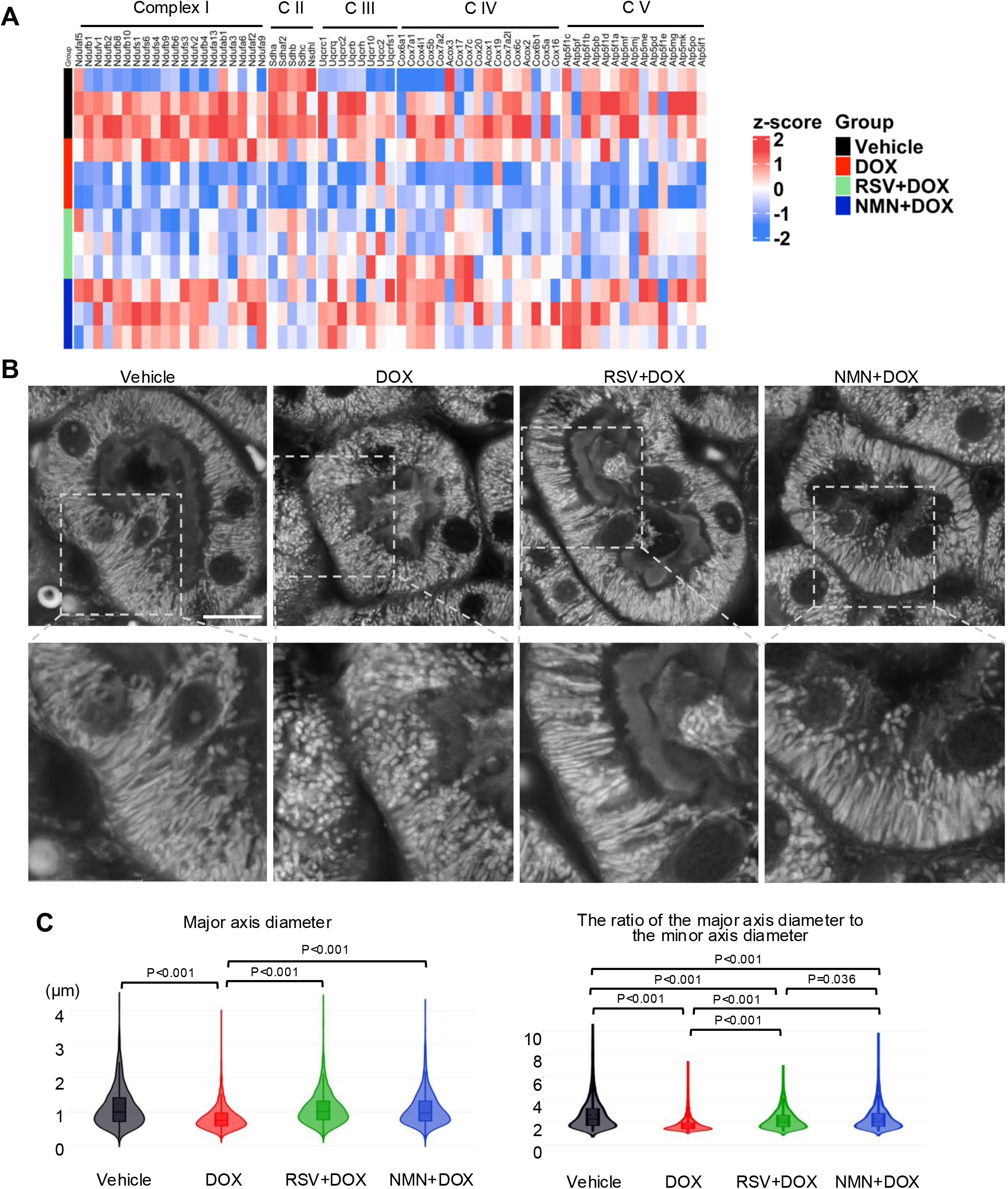
RSV and NMN ameliorate the DOX-induced mitochondrial structural abnormalities. (A) Heatmap of mitochondrial proteins belonging to the electron transfer system (complex I–V) from proteome analysis. Heatmap intensity indicates the level of gene expression (Z-score). (B) Representative structured illumination microscopy (SIM) images of mitochondria in proximal tubules from Masson’s trichrome-stained kidney sections. Scale bar, 5 μm. (C) Quantification of mitochondrial morphology, including major-axis length (left) and aspect ratio (major/minor axis; right). A total of 2,455, 3,386, 2,730, and 2,330 mitochondria were analyzed in the Vehicle, DOX, RSV+DOX, and NMN+DOX groups, respectively. Four proximal tubules from four mice per group were analyzed. Statistical analysis was performed using a one-way ANOVA on ranks, followed by Dunn’s multiple comparison test.

Quantitative morphometric analysis was performed by measuring mitochondrial area and the aspect ratio (major-to-minor axis length). Chronic DOX administration significantly reduced both major axis diameter and aspect ratio, indicating enhanced mitochondrial fragmentation in tubular epithelial cells. Importantly, co-treatment with RSV and NMN significantly attenuated these DOX-induced morphological alterations (Fig. 5B, C). Collectively, these findings demonstrate that chronic DOX exposure induces mitochondrial fragmentation in renal tubules and that pharmacological activation of the NAD⁺–Sirtuin axis mitigates these structural abnormalities.

### 3.7. *Ex Vivo* Seahorse Assay Demonstrates Impaired Mitochondrial Respiration Following Chronic DOX Exposure

To directly evaluate mitochondrial function in renal tubules, we employed a recently established *ex vivo* mitochondrial respiration assay with freshly isolated renal tubules, enabling assessment of mitochondrial bioenergetics under conditions that closely preserve in vivo physiology (Hammoud et al.). Mice were treated with the vehicle or DOX for four weeks, after which the renal tubules were immediately isolated and subjected to Seahorse metabolic flux analysis (Fig. 6A, B). Robust mitochondrial respiration and maximum OCR were observed in the vehicle-treated group. In contrast, renal tubules from DOX-treated mice exhibited a significant reduction in mitochondrial respiratory capacity, including a decrease in maximal respiration (Fig. 6C, D).

**Figure 6.**
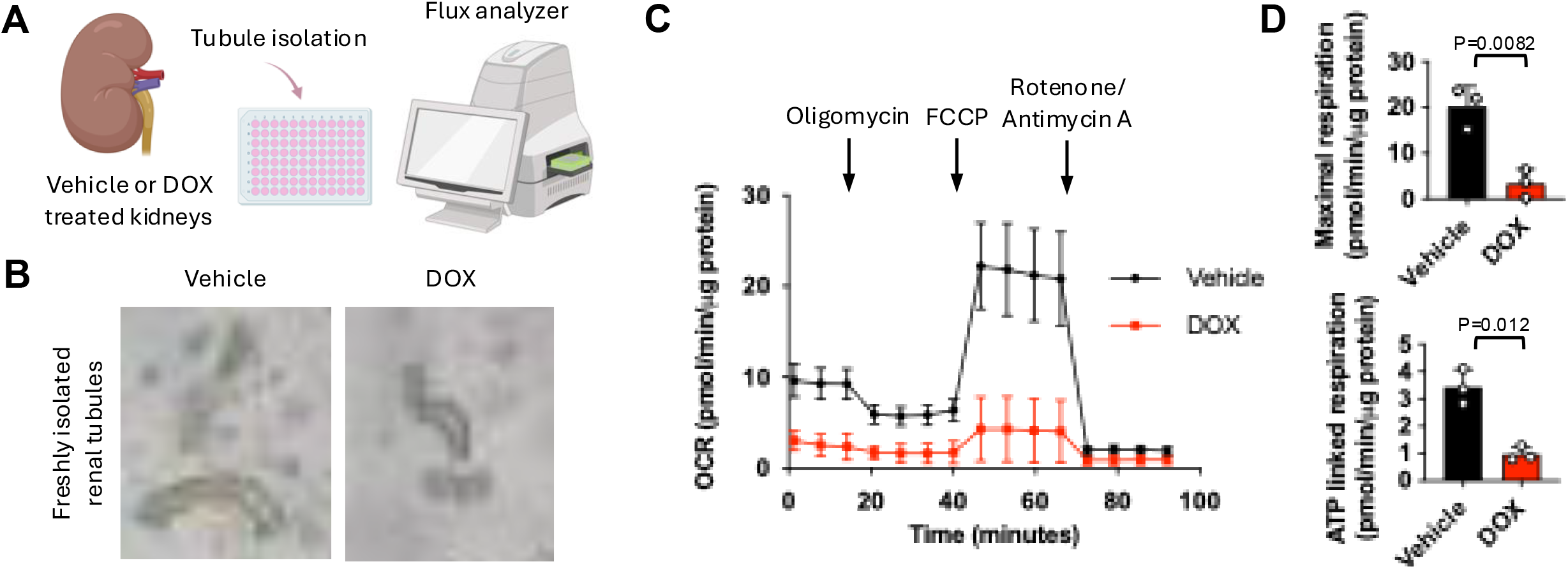
Chronic DOX treatment impairs mitochondrial respiration in freshly isolated renal tubules. **(A)** Schematic overview of *ex vivo* Seahorse analysis using freshly isolated renal tubules from Vehicle- or DOX-treated mice. Renal tubules were isolated from mouse kidneys and subjected to mitochondrial respiration analysis using the Seahorse XFe96 platform. **(B)** Representative images of freshly isolated renal tubules prepared from Vehicle-and DOX-treated mice. **(C)** Representative oxygen consumption rate (OCR) profiles of freshly isolated renal tubules. OCR was measured at baseline and after sequential injections of oligomycin, FCCP, and rotenone/antimycin A. **(D)** Quantification of maximal respiration and ATP-linked respiration. ATP-linked respiration was calculated by subtracting oligomycin-insensitive OCR from basal OCR, whereas maximal respiration was calculated by subtracting non-mitochondrial OCR, measured after rotenone/antimycin A injection, from carbonyl cyanide p-trifluoromethoxyphenylhydrazone (FCCP)-stimulated OCR. DOX-treated renal tubules exhibited significantly reduced mitochondrial respiratory capacity compared with vehicle-treated controls. Data are shown as mean ± SE. *P*-values are indicated in the graphs.

These findings confirm that chronic DOX exposure not only induces mitochondrial fragmentation but also impairs mitochondrial oxidative respiration in renal tubular cells.

### 3.8. RSV and NMN Attenuate Inflammatory Responses in Chronic DOX Nephropathy

Chronic inflammation is recognized as a key driver of tissue remodeling and fibrosis in both cardiotoxicity and CKD. In DOX-induced cardiomyopathy, injured cardiomyocytes trigger local inflammatory responses that contribute to progressive fibrosis (Shi et al. 2023). Similarly, persistent tubular injury has been implicated in chronic renal inflammation and CKD progression (Ide et al. 2021). Consistent with this concept, RNA sequencing analysis revealed significant enrichment of inflammatory pathways, such as adaptive immune response, response to interferon-beta, and leukocyte-mediated immunity, in the DOX-treated group compared with Vehicle controls (Fig. 7A).

**Figure 7.**
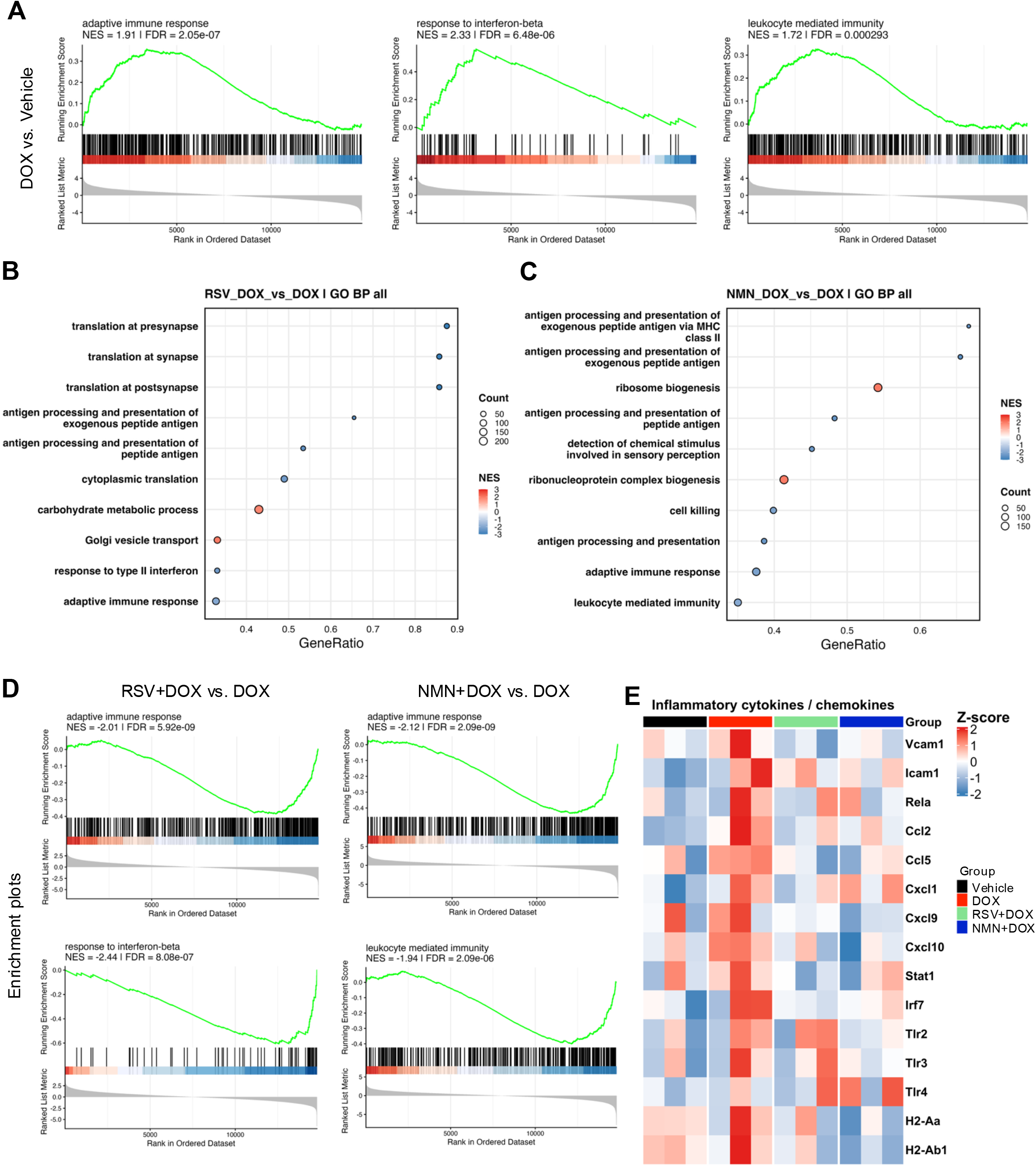
NMN and RSV suppress chronic DOX treatment-induced inflammatory responses. (A) Representative enrichment plots from RNA-seq data comparing the DOX and Vehicle groups. (B) Gene set enrichment analysis (GSEA) of RNA-seq data comparing the RSV+DOX and DOX groups. (C) GSEA of RNA-seq data comparing the NMN+DOX and DOX groups. (D) Representative enrichment plots from RNA-seq data comparing the NMN+DOX and DOX groups. (E) Heatmap showing the expression profiles of inflammatory cytokines, chemokines, and inflammation-related genes across experimental groups.

Importantly, a comparative GSEA between the DOX and RSV+ DOX or NMN+DOX groups showed the suppression of these inflammatory pathways in mice receiving sirtuin-activating treatments (Fig. 7B–D). Furthermore, the expression analysis of representative pro-inflammatory cytokines confirmed that DOX significantly increased inflammatory gene expression, whereas co-treatment with RSV or NMN attenuated this induction (Fig. 7E).

Collectively, these findings suggest that mitochondrial injury in tubular epithelial cells is accompanied by activation of inflammatory programs, and that pharmacological activation of the NAD⁺–Sirtuin axis mitigates chronic renal inflammation in this model.

## Discussion and Conclusions

In this study, we performed an integrated multi-omics analysis in a chronic DOX administration model for the first time and, in an unbiased manner, identified renal tubular mitochondrial dysfunction as a central mechanism of DOX-induced kidney injury. We further validated mitochondrial impairment using complementary structural and functional approaches and evaluated the therapeutic potential of activation of the NAD⁺–Sirtuin axis, previously implicated in DOX cardiomyopathy, in the context of renal tubular injury.

Our multi-omics analyses consistently demonstrated that chronic DOX exposure suppresses mitochondrial metabolic pathways in renal tissues. Clinically, there are patients in whom conventional markers, such as serum creatinine or overt proteinuria, remain within normal ranges, yet renal function gradually declines. Our findings suggest that subclinical tubular mitochondrial injury may precede measurable declines in conventional renal function markers, thereby contributing to the delayed onset of clinically apparent nephrotoxicity. Indeed, acute-phase evaluations demonstrate that immediately after administration of agents such as daunorubicin, urinary N-acetyl-β-D-glucosaminidase activity increases significantly, even when serum creatinine remains stable, serving as a sensitive indicator of tubular damage that precedes overt renal dysfunction (Bárdi et al. 2007). Anthracycline treatment, particularly at cumulative doses of ≥ 250 mg/m², has been reported as an independent risk factor for late-onset kidney failure in childhood cancer survivors (Dieffenbach et al. 2021). These findings suggest that mitochondrial abnormalities may occur prior to overt tubular injury and the subsequent decline in renal function, highlighting the potential utility of mitochondrial assessment as an early biomarker for detecting subclinical nephrotoxicity and predicting later renal dysfunction.

Mitochondrial injury was assessed by two complementary methods. First, SIM applied to Masson’s trichrome-stained sections enabled high-resolution visualization of mitochondrial morphology (Matsumoto et al. 2021). Importantly, unlike electron microscopy, this method does not require specialized sample preparation and can be readily applied to routinely processed Masson’s trichrome-stained paraffin sections, highlighting its potential translational utility for the clinical assessment of mitochondrial injury. Second, *ex vivo* Seahorse analysis of freshly isolated renal tubules allowed the functional evaluation of mitochondrial respiration while largely preserving the metabolic characteristics of native renal tubules. Consistent with our approach, recent studies using seahorse assays in freshly isolated renal tubules have demonstrated that this method enables the evaluation of tubular cell metabolism under conditions that more closely reflect the in vivo physiological state than conventional cultured cell systems (Hammoud et al. 2025). Furthermore, our findings indicate that this approach is well-suited for assessing drug-induced tubular injury and mitochondrial dysfunction, particularly in the setting of chronic DOX exposure. Importantly, the *ex vivo* metabolic profiling of isolated renal tubules may provide a sensitive platform for detecting early mitochondrial abnormalities associated with nephrotoxic injury.

Historically, DOX-induced organ toxicity has been predominantly studied in the heart (Wu et al. 2024; Li et al. 2019; Kuno et al. 2023; Wallace, Sardão, and Oliveira 2020). Both cardiomyocytes and renal tubular epithelial cells are highly enriched in mitochondria and rely heavily on oxidative metabolism, suggesting a shared vulnerability to mitochondrial insults (Doke and Susztak 2022; Brown et al. 2017). Sirtuin activation has previously been reported to ameliorate DOX cardiomyopathy in experimental models (Cheung et al. 2015; Tomczyk et al. 2022; Kuno et al. 2023). Extending these observations to the kidney, we found that pharmacological activation of the NAD⁺–Sirtuin axis markedly attenuated DOX-induced tubular injury and restored mitochondrial homeostasis. RSV is known to activate SIRT1, whereas NMN increases intracellular NAD^+^ levels, thereby activating multiple sirtuin family members, in addition to SIRT1. Previous studies suggested that increased NAD^+^ levels indirectly enhance SIRT3 activity. In our study, both RSV and NMN reduced acetyl-SOD2 levels, suggesting restoration of sirtuin activity. These findings suggest that activation of the NAD^+^–Sirtuin axis may exert renoprotective effects against DOX-induced mitochondrial injury and highlight SIRT3 as a potentially important therapeutic target. Further studies are warranted to clarify the relative contributions of SIRT1 and SIRT3 to renal protection.

This study had several limitations. First, it was conducted using a murine model, and extrapolation to human diseases requires cautious validation. Future studies that apply mitochondrial morphometric analysis to human renal biopsy specimens are important for determining their clinical relevance. Second, the potential contribution of cardiorenal interactions cannot be entirely ruled out. Tubule-specific genetic models, including conditional SIRT1 or SIRT3 knockouts or activation in renal tubular cells, are necessary to elucidate the cell-autonomous effects of sirtuin signaling in DOX-induced nephropathy. In addition, our findings suggest that both the heart and kidneys are protected through shared underlying mechanisms, particularly those involving mitochondrial dysfunction and the sirtuin pathway. These findings support a cardiorenal framework in which anthracycline-induced cardiac and renal toxicities share common mitochondrial mechanisms and may benefit from integrated therapeutic strategies. Such an integrated strategy may provide a novel therapeutic approach for the prevention and treatment of chemotherapy-induced multi-organ injuries.

In summary, our study provides the first unbiased multi-omics evidence that chronic DOX exposure induces mitochondrial dysfunction in renal tubules and demonstrates that pharmacological activation of the NAD⁺–Sirtuin axis confers tubular protection. These findings establish renal tubular mitochondrial dysfunction as a key mechanism underlying chronic anthracycline nephrotoxicity and identify the NAD⁺–Sirtuin axis as a promising therapeutic target for preventing chemotherapy-associated kidney injury.

## Author Contributions

K. Saito, A. Kuno, and K. Abe conceived and designed the study; K. Saito, T. Hori, R. Hosoda, Y. Saga, R. Numazawa, I. Nojima, K. T. Sato, Abe K, and A. Kuno performed the experiments and acquired the data; K. Saito, T. Hori, R. Hosoda, Y. Saga, R. Numazawa, I. Nojima, Y Tatekoshi, K. T. Sato, Abe K, and A. Kuno analyzed and interpreted the data. All the authors were involved in drafting and revising the manuscript. All the authors have read and approved the final version of the manuscript.

## Acknowledgements

This work was supported by the Northern Advancement Center for Science & Technology (NOASTEC) Foundation projects S-2-9 and H-1-6. The funders played no role in the study design, data collection, data analysis, decision to publish, or manuscript preparation.

## Conflict of Interest Statement

The authors declare no conflict of interest.

## Notes

### Competing Interest Statement

The authors have declared no competing interest.

